# Prophage-encoded hotspots of bacterial immune systems

**DOI:** 10.1101/2021.01.21.427644

**Authors:** François Rousset, Julien Dowding, Aude Bernheim, Eduardo P.C. Rocha, David Bikard

## Abstract

The arms race between bacteria and phages led to the emergence of a variety of genetic systems used by bacteria to defend against viral infection, some of which were repurposed as powerful biotechnological tools. While numerous defense systems have been identified in genomic regions termed defense islands, it is believed that many more remain to be discovered. Here, we show that P2- like prophages and their P4-like satellites have genomic hotspots that represent a significant source of novel anti-phage systems. We validate the defense activity of 14 systems spanning various protein domains and describe PARIS, an abortive infection system triggered by a phage-encoded anti-restriction protein. Immunity hotspots are present across prophages of distant bacterial species, highlighting their biological importance in the competition between bacteria and phages.

## Introduction

Bacteria employ an arsenal of defense strategies to overcome infection by bacteriophages (1). Recent studies have shown that bacterial immunity is much more diverse that previously envisioned, with newly uncovered defense systems spanning various mechanisms such as DNA restriction (2–5), abortive infection (6–12), but also chemical defense (13, 14). A series of remarkable discoveries stemmed from the observation that bacterial defense systems tend to cluster in genomic regions termed defense islands (15, 16). The systematic investigation of genes found in association with known defense genes has considerably expanded our knowledge of bacterial immunity (17, 18). Despite these remarkable findings, it is believed that many defense systems remain to be discovered.

Temperate phages can be stably maintained in their host cell as prophages for generations. During lysogeny, phage survival is tied to host survival, providing a selective pressure for temperate phages to carry “moron” genes that are not essential for the phage but enhance the fitness of their host (19). Some prophages encode toxins and virulence factors that increase the fitness of pathogenic bacteria during infection (20), while others promote bacterial growth in certain environments and resistance to stresses including antibiotics (21–23). Likewise, prophage-encoded defense systems indirectly enhance phage fitness by benefiting the host. Prophages have long been known to block infection by related or distant phages. The genetic control of lysogeny, which is maintained through the expression of a repressor of the lytic genes, ensures that incoming phages that share the same repressor will not enter the lytic cycle. In addition to repressor-based immunity, prophage-encoded systems can provide phage resistance through diverse mechanisms including modification of cell surface receptors (24), inhibition of DNA translocation into the cytoplasm (25), premature transcription termination (26) or abortive infection (9). A well-known example is that of the rex genes of bacteriophage λ which provide resistance against P22, T1, T5, T4 and T7 by triggering cell death after infection (27). More recent work has highlighted how prophages and phage satellites can mobilize defense systems in the context of inter-viral competition (28–32).

Phages have evolved anti-defense strategies in response to defense systems. The resulting arms race is driving genetic innovation and produced a stunning diversity of phage defense strategies and anti-defenses. As a consequence, there is a constant turnover of defense systems in bacteria and counter-defense systems in phages (*1*). While bacteria can readily accumulate diverse defense systems in their genome within defense islands, the genome of bacteriophages is typically constrained by the size of the DNA that can be packaged into the capsid, limiting the number of defense systems they can carry. Genetic diversity in a specific phage clade is typically constrained to specific regions of the phage genome. As an example, the enterobacteria P2-like phages harbor a large set of different genes at the Z/fun locus (*33*) thought to affect virulence. Anti-defense genes have also been found in diversity hotspots in phage genomes (*34, 35*).

Here, we show that hotspots of genetic diversity within the P2-like and P4-like prophage families constitute a large reservoir of bacterial immune systems, including known as well as many unknown defense systems. We describe in more details one of these systems, PARIS (<u>p</u>hage <u>a</u>nti-restriction-induced <u>s</u>ystem), an abortive infection system that triggers growth arrest upon sensing an anti-restriction protein. We also expanded our analysis and identified two more hotspots in prophages from Vibrionales and Bacillales. These findings highlight how prophages constitute a reservoir of defense proteins and point out to a new strategy to uncover novel bacterial immune systems.

## Results

### P4-like satellite phages encode a hotspot for bacterial immune systems

We recently identified a retron in a P4-like satellite prophage that is responsible for the essentiality of the exodeoxyribonuclease I SbcB in the *E. coli* strain H120 (*36*). Retrons are intriguing genetic elements that comprise a reverse-transcriptase (RT) and a multi-copy single-stranded DNA and were recently shown to form tripartite toxin/antitoxin (TA) systems involved in anti-phage defense (*7, 8, 18, 37*). Since P4-like prophages are very prevalent in Enterobacteria (including in around a third of *E. coli* isolates (*38*)), we inspected other *E. coli* strains and noticed that each P4-like element carried a different genetic element at the same locus between the polarity suppression protein (*psu*) and the integrase (*int*) (**Fig. 1A**). These elements are adjacent to the *cos* packaging signal and found in either orientation. They include a variety of uncharacterized proteins as well as known defense proteins belonging to type II and III restriction-modification (RM) and retrons, a few of which had previously been reported (*39, 40*). We identified 5,251 occurrences of this locus in >20,000 *E. coli* genomes (**Table S1, see Methods**), together encoding >300 different gene arrangements. We analyzed 121 arrangements occurring at least five times (together accounting for 94.4% of all loci) (**Fig. S1**) and identified 81 different genetic systems (**Fig. 1B, Table S2**). They can be broadly grouped into three categories : (i) known defense systems such as a type-III restriction enzyme, the type-II EcoO109I RM system (*39*), retrons (*7, 8, 18, 37*), Gabija (*17*) or Septu (*17*); (ii) uncharacterized systems with protein domains that were previously associated with bacterial immunity such as SIR2 (*17, 41*), TIR (*17*) and HEPN (*18, 42*); (iii) systems encoding proteins with no annotated domain or whose domain is currently not associated with anti-phage defense, such as NACHT and various domains of unknown function (DUFs). Based on these observations, we hypothesized that the P4-encoded locus between *psu* and *int* may constitute a reservoir of bacterial immune systems that likely participate in interviral competition.

**Figure 1.**
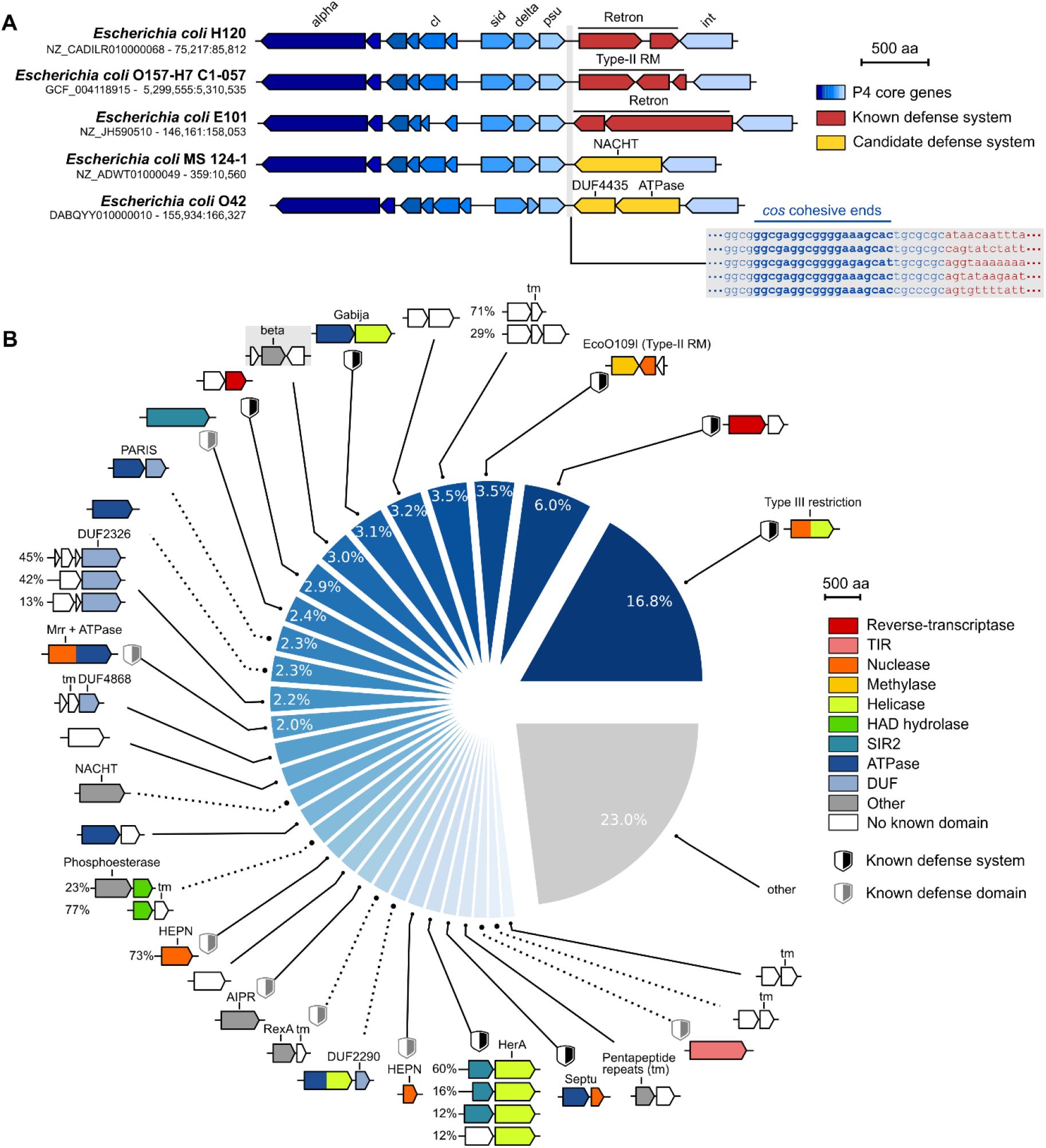
A diversity of genetic systems encoded on P4-like phages in *E. coli*. (**A**) Genomic visualization of P4-like prophages in five *E. coli* strains, highlighting genetic diversity between *psu* and *int* genes, including known anti-phage defense systems. Genome accession numbers and positions are shown on the left. The DNA sequence of the *cos*-proximal region in these strains is highlighted with conserved sequences in blue and variable sequences in red. (**B**) Systematic analysis of genetic systems encoded between *psu* and *int* identified in 26% (5,251/20,125) of analyzed *E. coli* genomes. The pie chart shows the proportion of loci encoding each system (considering 81 curated systems present in at least 5 occurrences, see Methods). The 30 most abundant systems are shown as gene cassettes colored by protein domains (not to scale). Among these, 8 validated systems providing phage defense (**Fig. 2A**) are highlighted with a dashed link. The system present in the P4 reference genome is highlighted with a grey background. When a system comprises accessory genes, different variants are shown with the percentage of each occurrence shown on the left. TIR: Toll/Interleukin-1 Receptor, HAD: haloacid dehydrogenase-like, SIR2: sirtuin, DUF: domain of unknown function, tm: transmembrane domain.

To test for anti-phage activity, we cloned 16 systems from *Escherichia* and *Klebsiella*, comprising each of the three above-mentioned categories. We introduced them into *E. coli* K-12 MG1655 and challenged each resulting strain with an array of 9 coliphages spanning most common phage families (**Table S3**). When compared to a control vector encoding a green fluorescent protein (GFP), most systems (68.8%; 11/16) provided resistance to at least one phage (**Fig. 2A, Fig. S2 and Table S4**). This suggests that most of the diverse genetic systems encoded at this hotspot are involved in anti-phage defense. A retron from *E. coli* E101 associated with a transmembrane effector protected against P1vir and 186cIts, as expected from previous findings (*7, 8, 18, 37*). Among unknown systems with known defense domains, we identified a HEPN-family protein as well as other single-protein systems that comprise either an AAA+ ATPase, a sirtuin (SIR2) or a toll/interleukin-1-receptor-like (TIR) domain. Various classes of ATPases from the AVAST systems have been recently described (*18*) while SIR2 and TIR domains are present in the Thoeris defense system (*12, 17*) and are important determinants of Eukaryotic immunity (*41, 43*).

**Figure 2.**
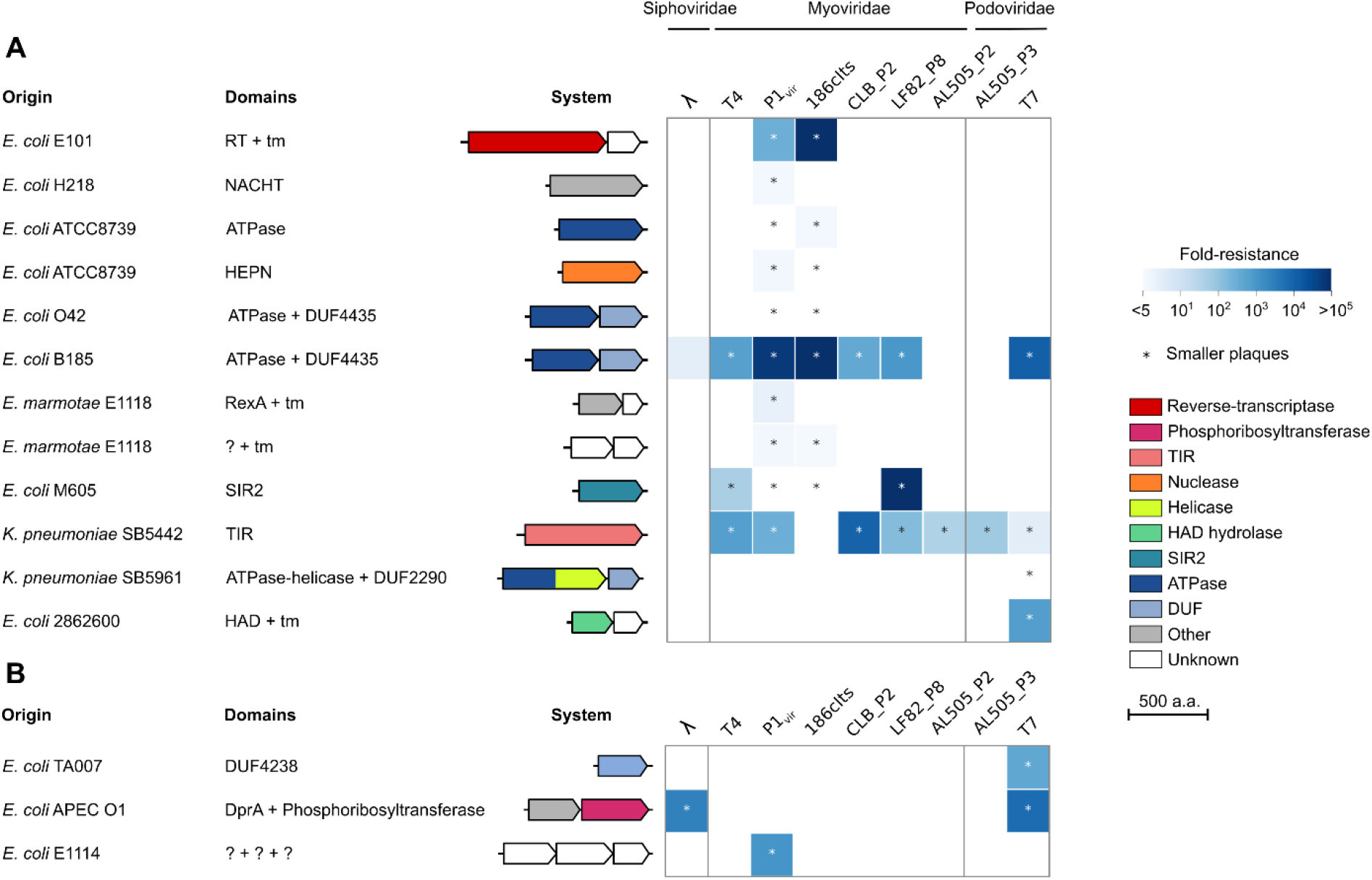
Prophage-located hotspots encode a variety of bacterial immune systems. Phage resistance heatmaps of the validated defense systems show the median fold resistance of three independent replicates against a panel of 9 phages (**Table S3**). We verified the anti-phage activity of (**A**) 12 occurrences of 11 systems from the P4-encoded hotspot, some of which are highlighted in Fig. 1B and (**B**) 3 systems from the P2-encoded hotspot. Genes are colored by protein family (not to scale). Genome accession numbers are provided in **Table S4**. RT: reverse-transcriptase, TIR: Toll/Interleukin-1 Receptor, HAD: haloacid dehydrogenase-like, SIR2: sirtuin, DUF: domain of unknown function, tm: transmembrane domain.

We also identified a distant analog of the RexAB system from phage λ (*27*) that provides moderate protection against P1_vir_. Systems that have never been linked to bacterial immunity include a protein with a NACHT NTPase domain that is involved in programmed cell death and immunity in Eukaryotes (*44*), as well as a two-protein system that comprises an unknown protein associated with a transmembrane effector. We also describe an ATPase associated with a DUF4435 protein and two systems that specifically inhibit the growth of phage T7: an ATP-dependent helicase associated with a DUF2290 domain and a HAD-like hydrolase associated with a transmembrane protein.

Taken together, our findings show that P4-like phages carry a hotspot that constitutes a reservoir of anti-phage defense systems. P4-like elements identified in other Enterobacteriaceae than *E. coli* also showed candidate defense genes at the same locus (**Fig. S3**).

### PARIS triggers growth arrest upon sensing an anti-restriction protein

We further investigated a new defense system from *E. coli* B185 comprising an AAA+ ATPase associated with a DUF4435 protein that provides a broad defense phenotype by protecting against 7/9 phages of our panel (**Fig. 2A**). We monitored the growth of *E. coli* K-12 MG1655 carrying this system or a control plasmid during infection by phage T7 at a low (0.005) or high (5) multiplicity of infection (MOI). In the presence of a control plasmid, infection led to population collapse regardless of the MOI (**Fig. 3A**). In contrast, the system provided partial resistance to T7 at a low MOI but led to a growth halt after infection at a high MOI. In addition, the number of infected cells that released viable phage was reduced by ∼15- fold (95% CI: [12.1;18.5]) in the presence of the system. This defense system therefore seems to arrest growth of infected cells, preventing the completion of the phage cycle, a typical feature of abortive infection systems (*45*). Deletion of either the ATPase or the DUF4435 gene abrogated defense (**Fig. S4**). We aimed at identifying the phage-encoded trigger by isolating phage escapers that were able to overcome this system. We successfully isolated T7 mutants that improved the EOP ∼10^5^-fold compared to wild-type T7 (**Fig. 3B**). Whole genome sequencing of 4 mutants revealed that all of them carried a mutation (T54V) in gene 0.3 which encodes Ocr, an anti-restriction protein that inhibits RM and BREX systems by mimicking the structure of DNA (*46, 47*). In the presence of the system, the transformation efficiency of an Ocr-expressing plasmid was ∼100-fold lower than in the absence of the system, while there was no difference when transforming a GFP-encoding plasmid as a control (**Fig. 3C**). This confirms the toxicity of Ocr in the presence of this defense system. We can therefore describe it as an anti-anti-restriction system, which we decided to name PARIS (<u>p</u>hage <u>a</u>nti-restriction-induced <u>s</u>ystem). This represents a remarkable evolutionary strategy where cells have evolved to use a counter-defense protein as a trigger for abortive infection, similarly to the Prr system (*48*). This strategy allows cells to undergo self-sacrifice only when an incoming phage can overcome the first line of defense (e.g. a RM or BREX system), therefore maximizing both cellular and population-level survival. We detected PARIS in more than 4% of bacterial and archaeal genomes from diverse clades (**Fig. S5**), either in a two-gene cassette as described here or in a single-gene fusion comprising both the ATPase and DUF4435 domains, a strong evidence of tight functional interaction (*49*).

**Figure 3.**
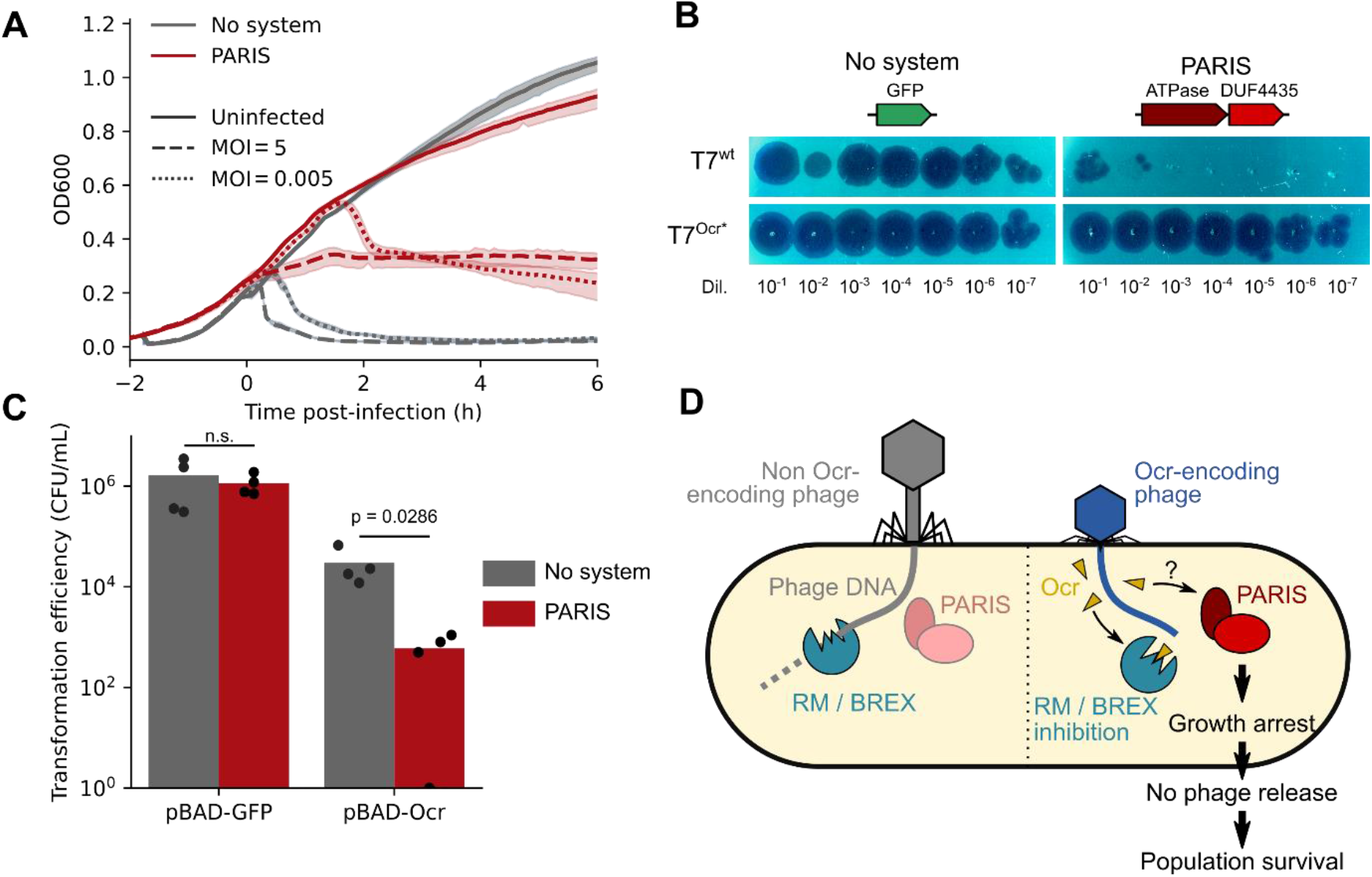
PARIS senses phage-encoded anti-restriction. (**A**) Time course experiment with cells harboring a control plasmid or a PARIS-encoding plasmid. Cells were kept uninfected or were infected with T7 at a high or low multiplicity of infection (MOI) once cells reached OD∼0.2. Each curve shows the mean of three technical replicates with the standard deviation shown as a transparent area. (**B**) Serial dilutions of a high titer lysate of wild-type or mutated (Ocr*) T7 phage on a lawn of cells harboring a control plasmid or a PARIS-encoding plasmid, representative of 3 independent replicates. (**C**) Transformation efficiency of a plasmid expressing a green fluorescent protein (GFP) or T7 Ocr protein. Bars show the mean of 4 independent replicates. The p-value of a two-sided Mann-Whitney test is shown. (**D**) Current model for the defense activity of PARIS.

### A hotspot in P2-like phages comprises new bacterial immune systems

P4 is a satellite phage that lacks structural genes encoding capsid and tail proteins. Instead, it hijacks capsids from helper phages of the P2 family and uses the P2 terminase to package its own genome into modified P2 capsids (*50*). As a consequence, P4 and P2 share a core *cos* packaging signal (*51*). The fact that the defense hotspot is directly adjacent to the *cos* site in P4-like phages prompted us to inspect the region adjacent to the *cos* site in P2-like phages, which is located between the replication protein gpA and the portal protein gpQ. The P2 reference genome (NC_001895) actually encodes the phage exclusion genes *old* and *tin* at this locus (*52*). We performed a systematic analysis of this locus in all *E. coli* genomes (**see Methods**). Our results show that this region is another hotspot for genetic diversity (**Fig. S6 and Table S5**), with 1,699 different gene arrangements detected from 18,150 occurrences of this locus. We curated the arrangements occurring at least ten times and identified 147 genetic systems comprising genes encoding known defense proteins such as retrons and type-III restriction enzymes, and other genes likely involved in other functions. For instance, more than a fourth of P2-like phages encode genes which likely belong to a plasmid partitioning system that could allow these prophages to be maintained as plasmids, while around 2% encode a cytolethal distending toxin which likely participates in bacterial virulence (*53*). Importantly, some systems are common between P4- and P2-encoded hotspots but are present in different frequencies, while many systems are found in one hotspot but not in the other. P2-like hotspots tend to be slightly larger than those of P4-like (3.2 vs 2.6 kb on average) (**Fig. S7**), likely reflecting the pressure faced by the latter to package their genome into a capsid of reduced size which can only accommodate ∼12kb (*54*). Accordingly, P2 and P4 encode mostly short systems and lack the large CRISPR-Cas systems and type-I RM that can be found in many bacterial genomes or in some large plasmids.

We cloned 8 more systems from the P2-encoded hotspot and tested their activity against the 9 phages described above. We observed defense activity for 3 systems (**Fig. 2B**). The first one encodes a protein with a DUF4238 domain (whose molecular function is currently unknown) that provides strong resistance to T7. The second system comprises three proteins with no known domain and protects against P1vir. The third system inhibits infection by phages T7 and λ and encodes a DNA-processing chain A protein (DprA) associated with a phosphoribosyltransferase (PRTase). DprA-like proteins are involved in DNA transformation in naturally competent bacterial species by binding to single-stranded DNA and interacting with RecA (*55*). In *Helicobacter pylori*, DprA also promotes DNA methylation and alleviates restriction barriers during natural transformation (*56*). Non-competent species also carry DprA-like proteins whose role has remained elusive. Our finding provides another function to DprA-like proteins in non-competent bacteria. DprA may bind and deliver phage DNA to the PRTase effector which may neutralize it. Alternatively, the frequent association of PRTase effectors with retrons (*7, 57*) suggests that the PRTase could trigger cell suicide after phage DNA sensing by DprA.

Taken together, our results suggest that both P4 and P2-like phages carry a hotspot of genetic diversity adjacent to the *cos* site and which includes a large diversity of genes involved in phage defense.

### Hotspots for bacterial immunity identified in other species

We wondered whether such hotspots also exist in prophages belonging to more distant bacterial species. We were able to identify at least two more occurrences of such hotspots through manual examination of loci that harbor some of the systems described here. The first hotspot is prevalent in *Vibrionaceae* between the cI repressor and the integrase of a P2-related phage that is similar to *Vibrio cholerae* phage K139 (*58*) and *Aeromonas* phage ΦO18P (*59*) (**Fig. 4A**). The second one is encoded on a prophage from *Bacillus* between a transcriptional regulator and a peptidase adjacent to the integrase (**Fig.4B**). In both cases, hotspots encode known defense systems such as RM, Septu as well as the ATPase + DUF2290 system and PARIS described above. While most proteins in these loci have currently not been linked to bacterial immunity, many of them have known defense domains such as nucleases, kinases and ATPases. This analysis reveals that prophage-encoded diversity hotspots are present in various bacterial organisms and likely constitute a significant reservoir of new bacterial immune systems across diverse bacterial phyla.

**Figure 4.**
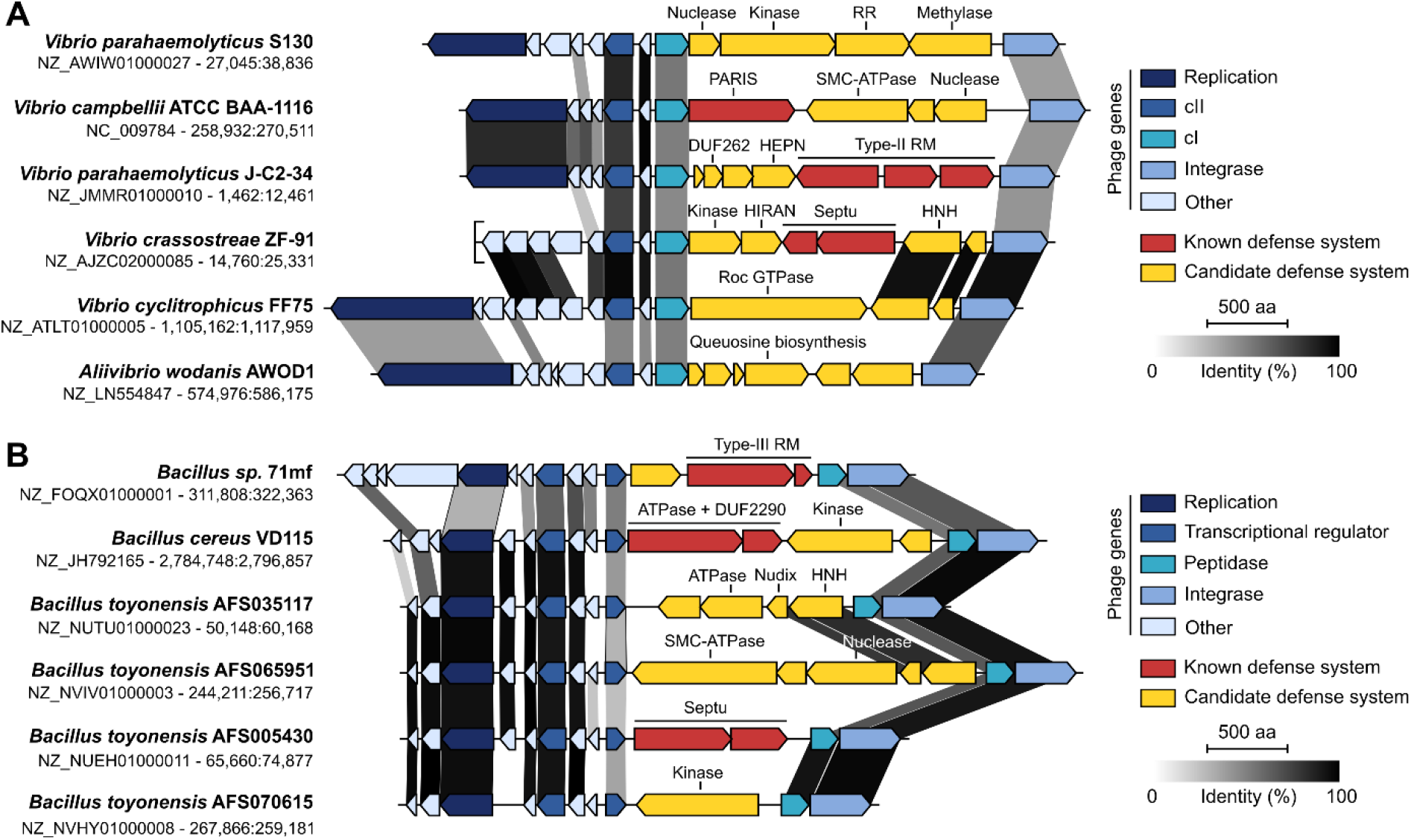
Hotspots for bacterial immune systems encoded on other prophage genomes. Genomic view of hotspots encoded on prophages from *Vibrionales* (**A**) and *Bacilliales* (**B**). Phage genes are shown with different shades of blue. Grey shades show the percentage of identity between homologous proteins from different genomes. Genome accession numbers and positions are shown on the left.

## Discussion

Our findings showcase a diversity of bacterial immune systems in prophage-encoded hotspots and uncover a new strategy to identify novel anti-phage systems. The different hotspots that we describe constitute a significant reservoir of putative immune systems, illustrating the vastness of what remains to be discovered. Importantly, the proportion of P4-encoded systems that showed anti-phage activity reached 69% (11/16) and would likely increase with the testing of more phages. The presence of such a diverse panel of immunity proteins illustrates the extent of inter-viral competition and likely mirrors the diversity of phages encountered by *E. coli* and the anti-defense strategies they deploy.

The acquisition by a temperate phage of a defense system for which other phages are sensitive favors the phage because it protects its lysogen. Yet, as the frequency of resistant phages rises in the population, the defense system becomes less and less adaptive. This eventually results in the selection of temperate phages that acquired novel defense systems for which the other phages are still sensitive. In this never-ending eco-evolutionary dynamic, it is interesting to consider the mechanisms yielding such a dramatic rate of gene exchange specifically at the hotspots. In both P2-like and P4-like phages, the hotspots are directly adjacent to the *cos* sequence. It is tempting to speculate that cohesive ends generated after cleavage by the terminase or made available immediately after injection of the phage DNA into the cytoplasm could provide a substrate for recombination upon coinfection by several P2-like or P4-like phages, or upon co-induction of resident prophages. A second recombination event mediated by homologous recombination would result in genetic exchange of the locus. Further experimental work will be necessary to validate this or other possible recombination pathways.

The integration of temperate phages, their satellites (this work), and integrative conjugative elements (*60, 61*) carrying defense systems provides a mechanism for the genesis and evolution of defense islands in bacterial genomes. These mobile elements tend to integrate at a small number of chromosomal hotspots (*62*). Their inactivation by mutation results in rapid gene loss until only the functions adaptive to the bacterial host remain (*63*). In fact, we have found occurrences of P4-like phages that are integrated near defense islands or integrative conjugative elements that carry defense genes (**Fig. S8**). The high turnover of mobile elements in a few chromosomal hotspots can lead to the accumulation of defense systems in specific loci. The bacterial genome is much less constrained than those of phages and can thus accumulate many more systems in the same hotspot. In this hypothesis, bacterial defense islands against mobile elements result from the accumulation of defense systems from mobile elements themselves. Our findings also provide further evidence of the genetic entanglement that exists between defense systems, viruses and mobile genetic elements in general (*32*).

We anticipate that the systematic search for anti-phage defense hotspots in genomic but also metagenomic datasets will uncover many novel immune systems. Unlike previous systematic approaches (*17, 18*), our method only requires short contigs and is as such applicable to metagenomic and virome data. Immunity factors carried by viruses have been described across all the other domains of life (*32*), and it is tempting to hypothesize that viral defense hotspots might exist there as well.

## Material and Methods

### Identification of prophage-encoded systems

We downloaded all 20,125 *E. coli* genomes and encoded protein sequences available on Genbank in August 2020 with any assembly status. We used the presence of the *psu* gene to pinpoint P4-like elements. To extract the systems located between *psu* and *int* genes in these elements, we performed a blastp search (version 2.9.0) against all *E. coli* proteins with an E-value threshold of 10_^-10^_ using the Psu protein sequence from *E. coli* E101 as a query (WP_000446153.1). For each hit, we searched for the presence of an integrase by keyword (“integrase”) in the 10 downstream genes and retrieved the genes located in between, yielding a total of 5,251 loci after discarding loci that were across two contigs (**Table S1**). To reconstruct systems and assess their frequency, all 10,860 proteins from all loci were then clustered using MMseqs2 (version 8ebc9d16b2679eb485803259c8280127801e074b) (*1*) using a coverage threshold of 60% (-c 0.6 option), resulting in 541 clusters. For each locus, we then defined protein arrangements as a suite of clusters. We identified a total of 318 different protein arrangements. We manually curated the 121 protein arrangements that were present at least 5 times (together accounting for 94.4% of all loci) and grouped together highly similar systems sharing a core set of genes, resulting in 81 systems summarized in **Table S2**.

To extract the systems located between gpA and gpQ genes in P2-like prophages, we performed a blastp search against all *E. coli* proteins with an E-value threshold of 10^−10^ using the gpQ protein from Enterophage P2 as a query (NP_046757.1). Similarly as described above, we searched for the presence of gpA by keyword (“replication endonuclease”) in the 10 downstream genes and retrieved the genes located in between as candidate loci, yielding a total of 18,150 loci after discarding loci that were across two contigs. All 66,498 proteins from all loci were then clustered with MMseqs2 (-c 0.6 option), resulting in 846 protein clusters, and protein arrangements were reconstructed as described above, to the exception that we also discarded arrangements that only comprised proteins shorter than 120 amino acids. We identified a total of 1699 different arrangements (**Table S5**), including 316 present in at least 10 occurrences (together accounting for 82% of all loci) that were manually curated as described above to yield 147 different systems.

Selected loci were visualized using clinker and clustermap.js (*2*).

### Detection of PARIS

In order to detect two-protein occurrences of the PARIS system (ATPase + DUF4435), we downloaded HMM profiles for the AAA_15 (PF13175), AAA_21 (PF13304) and DUF4435 (PF14491) protein families from the pfam database (3). We then used these profiles to detect PARIS with MacSyFinder (v. 1.0.2) (*5*), requiring the two genes to be present (either AAA domain AND DUF4435). To detect single protein fusions, we browsed the “domain organisation” page of the DUF4435 pfam to look for proteins having both AAA_15/AAA_21 and DUF4435 domains. We selected 10 protein sequences with AAA_15+DUF4435 domains and 10 protein sequences with AAA_21+DUF4435 domains and built a single HMM profile from these. We then used this profile to detect PARIS with MacSyFinder (v. 1.0.2) (*5*) in default mode with the parameter “loner”. Using these detection rules, we analyzed 13,512 complete genomes retrieved from NCBI RefSeq in May 2019, representing 4,010 and 220 species of Bacteria and Archaea respectively.

### Strains and media

We used *E. coli* K-12 MG1655 to clone and test candidate defense systems as well as for subsequent experiments. Lysogeny broth (LB) was used as a liquid medium and LB supplemented with 1.5% agar was used as a solid medium. Kanamycin (Kan, Sigma) and carbenicillin (Carb, Euromedex) were used at 50 μg / mL and 100 μg / mL respectively.

### Cloning candidate defense systems

Systems were amplified from source strains, to the exception of 3 systems which were synthesized (Twist Bioscience): the active system from *E. coli* 2862600 as well as two systems from *E. coli* HVH3 and O157:H7 FRIK944 for which we did not detect any activity against our phage panel. All systems were amplified with their native promoters by Phusion PCR (ThermoFischer) using primers listed in **Table S4**. We used as vector pFR66, a low-copy plasmid with a pSC101 origin of replication, a kanamycin resistance cassette and a superfolder GFP gene (DNA sequence is provided in **Table S6**). pFR66 was made linear by PCR with primers 5’-TTTTGCCTCCTAACTAGGTC-3’ and 5’-CCAGGCATCAAATAAAACGAAAGGCTCAGT-CGAAAGAC-3’. The GFP gene was replaced by each candidate system using the Gibson method (*6*). All systems were transformed into electrocompetent *E. coli* K-12 MG1655 cells which were prepared as follows: following 100-fold dilution of an overnight culture in 200 mL of LB, cells were grown to OD ∼1, spun (4,000 g w 7 min) and washed three times in ice cold water, before concentration in ∼300 µL of 10% glycerol. One microliter of dialyzed Gibson assembly product was transformed into 20 µL of cells. After 1h of recovery at 37°C in LB medium, transformants were selected on LB + Kan plates. All constructions were verified using Sanger sequencing.

### Phage plaque assays

*E. coli* K-12 MG1655 strains carrying each of the systems or the control plasmid pFR66 were grown overnight in LB + Kan. Bacterial lawns were prepared by mixing 250 µL of a stationary culture with 62.5 µL of CaCl_2_ 1M and 12.5 mL of LB + 0.5% agar and the mixture was poured onto large square plates (12×12 cm) of LB + Kan. Serial dilutions of high-titer (>10^8^ pfu/mL) stocks of phages λ, T4, P1vir (a lytic-only variant of P1), 186cIts (a thermosensitive variant of 186), CLB_P2 (*7*), LF82_P8 (*8*), AL505_P2 (*9*) and AL505_P3 (*9*) were spotted on each plate and incubated overnight at 37°C. For phage T7, plates were incubated overnight at room temperature. Information related to our phage panel is provided in **Table S3**. The next day, plaques were counted and the fold resistance was measured as the number of plaques in the control plate divided by the number of plaques in the presence of each system. When plaques were too small to be counted individually, we considered the most concentrated dilution where no plaque was visible as having a single plaque. The complete list of validated systems is provided in **Table S4**.

### Time course infection experiments

Overnight cultures of K-12 MG1655 cells harboring a control plasmid or a plasmid encoding PARIS from *E. coli* B185 were diluted to OD ∼ 0.04 in LB + Kan and arrayed in a 96-well plate. Growth was then monitored in three replicates every 5 min on an Infinite M200Pro (Tecan) at 37°C with shaking. When OD reached ∼0.3, cells were either kept uninfected or infected with ∼2.10^8^ pfus (MOI ∼ 5) or ∼2.10^5^ pfus (MOI ∼ 0.005) of phage T7. Growth was then monitored for 6 h post-infection.

### Efficiency of centers of infection

To measure the number of infective centers, cells carrying PARIS or a control plasmid (pFR66) were grown in LB + Kan at 37°C until OD∼0.4. Cells were infected with T7 phage at a multiplicity of ∼0.01 and incubated at 37°C for 5 min. Cells were then centrifuged (6,000 g – 3 min) and resuspended in LB + Kan to eliminate free phage, and serial dilutions in 100 µL of LB + Kan were prepared. To each dilution, we added 100 µL of phage-sensitive cells (MG1655 + pFR66), 5 mL of LB + 0.5% agar and CaCl_2_ (5mM final concentration) and the mix was poured onto a LB + Kan plate. Care was taken to make sure the experiment was finished before lysis of infected cells (∼20 min post-infection). The next day, infective centers were measured as the number of plaque-forming units on each plate.

### Isolation and sequencing of mutant phages

We isolated T7 mutants that overcome PARIS by picking plaques in the lowest dilutions of a spot assay on a lawn of cells carrying PARIS from *E. coli* B185. For each putative mutant, plaques were reisolated on a lawn of K-12 MG1655 cells. The resistance phenotype of each mutant was then verified by comparing the number of plaques in the presence or absence of PARIS as shown in **Fig. 2**. To extract phage DNA, 500 µL of high titer stocks (∼10^10^ pfu/mL) of wild-type of mutant T7 were treated with TURBO DNAse (Thermo Fisher Scientific) for 30 min at 37°C. DNAse was inactivated with the addition of 5 µL of EDTA 0.5 mM and 25µL of inactivation reagent at 65°C for 10 min. The supernatant was then treated with 0.5 mg/mL of proteinase K (Eurobio) and SDS 0.5% to release phage DNA from capsids. DNA was then purified as follows: 500 µL of a phenol-chloroform-isoamylalcohol (PCI, 25:24:1) solution (Sigma) were added to the sample which was then vortexed and centrifuged (6,000 g – 5 min). To 500 µL of the upper aqueous phase were added another 500 µL of PCI solution. After vortexing and centrifugation (6,000 g – 5 min), the upper aqueous phase was transferred to a tube containing 500 µL of chloroform. The sample was further vortexed and centrifuged (6,000 g – 5 min), and the upper aqueous phase was transferred to a tube containing 500 µL of cold isopropanol and incubated for 2h at −20°C to precipitate DNA. After centrifugation (16,000 g – 1 min), the DNA pellet was washed with 75% ethanol. Finally, the pellet was air-dried and resuspended in 50 µL of distilled water. Next-generation sequencing was performed using a Nextera XT DNA library preparation kit and the NextSeq 500 sequencing systems (Illumina) at the Mutualized Platform for Microbiology (P2M) at Institut Pasteur. Mutations were identified by mapping sequencing reads to the T7 reference genome using breseq (v. 0.33.2).

### Transformation assays

To verify the involvement of Ocr in the activation of PARIS, we cloned either a GFP or the T7 Ocr protein on the pBAD18 vector as follows: using Phusion PCR (ThermoFischer), a GFP fragment was amplified using primers 5’-ACCCGTTTTTTTGGGCTAGCGAATTGATATCCGGAGGCATATCAA-3’ and 5’-GCCTTTCGTTTTATTTG- ATGCCTGGTTTGTAGAGTTCATCCATGC-3’ while Ocr was amplified from T7 genome using primers 5’- GCCTTTCGTTTTATTTGATGCCTGGTTACTCTTCATCCTCCTCGTACTCC-3’ and 5’-ACCCGTTTTTTTGGGCTAGCG- AATTGCAAGGTGCCCTTTATGATA-3’. The pBAD18 backbone was amplified using primers 5’- AATTCGCTAGCCCAAAAAAACGG-3’ and 5’-CCAGGCATCAAATAAAACGAAAGGCTCAGTCGAAAGAC-3’. Inserts were cloned into the backbone using the Gibson method (*6*). Constructions were electroporated into K-12 MG1655 cells and validated by Sanger sequencing, before plasmid extraction by miniprep (Macherey-Nagel). Electrocompetent cells of MG1655 carrying PARIS from *E. coli* B185 or a control plasmid were prepared as described above. A hundred nanograms of plasmids were electroporated in both cell types and incubated for 1h at 37°C for recovery. Serial dilutions were then plated on LB agar plates supplemented with 50 µg/mL of kanamycin, 100 µg/mL of carbenicillin and 0.3% of arabinose and CFUs were counted the next day.

### Identification of hotspots in other bacterial species

From the IMG database (*10*) (accessed on December 1^st^ 2020), we ran the built-in blastp function (e-value 10^−10^) using the DUF2290 and DUF4435 proteins from *K. pneumoniae* SB5961 and *E. coli* B185 respectively as queries. We then inspected the genomic neighborhood of distant hits using the online interface. We identified candidate hotspots when a neighboring gene frequently occurred next to the DUF2290 or DUF4435 proteins. In this way, we identified the cI gene from *Vibrionales* shown in **Fig. 4A** as well as the peptidase gene from *Baciliales* shown in **Fig. 4B** as potential locations for hotspots. To verify this, we then performed a new blastp search using these genes as queries against the IMG database and inspected the genomic neighborhood of their loci.

## Acknowledgments

We thank Erick Denamur for providing *E. coli* strains and Sylvain Brisse for providing *K. pneumoniae* strains. We also thank Laurent Debarbieux and Mathieu de Jode for providing phages CLB_P2, LF82_P8, AL505_P2 and AL505_P3, the P2M platform (Institut Pasteur, Paris, France) for genome sequencing, Rémi Denise for his assistance on Fig. S5, and Alicia Calvo-Villamañán and Raphaël Laurenceau for their contribution to the PARIS acronym. F.R. is supported by a doctoral fellowship from Ecole Normale Supérieure. This work was supported by the European Research Council (ERC) under the Europe Union’s Horizon 2020 research and innovation program (grant agreement No [677823]), by the French Government’s Investissement d’Avenir program and by Laboratoire d’Excellence ‘Integrative Biology of Emerging Infectious Diseases’ (ANR-10-LABX-62-IBEID), F.R. is supported by a doctoral scholarship from Ecole Normale Supérieure. E.R. was partially supported by the “Fondation pour la Recherche Médicale” (Equipe FRM EQU201903007835).

## Author contributions

F.R. and D.B. conceived the project. F.R. and A.B. analyzed the data. F.R. and J.D. performed experiments. F.R, A.B., E.R. and D.B. interpreted the results and wrote the manuscript.

## Data and materials availability

All data is available in the main text or the supplementary materials.

## References

1. A. Bernheim, R. Sorek, The pan-immune system of bacteria: antiviral defence as a community resource.Nat. Rev. Microbiol. 18 (2020), pp. 113–119.

2. T. Goldfarb, H. Sberro, E. Weinstock, O. Cohen, S. Doron, Y. Charpak-Amikam, S. Afik, G. Ofir, R. Sorek, BREX is a novel phage resistance system widespread in microbial genomes. EMBO J. 34, 169–183 (2015).

3. G. Ofir, S. Melamed, H. Sberro, Z. Mukamel, S. Silverman, G. Yaakov, S. Doron, R. Sorek, DISARM is a widespread bacterial defence system with broad anti-phage activities. Nat. Microbiol. 3, 90–98 (2018).

4. X. Xiong, G. Wu, Y. Wei, L. Liu, Y. Zhang, R. Su, X. Jiang, M. Li, H. Gao, X. Tian, Y. Zhang, L. Hu, S. Chen, Y. Tang, S. Jiang, R. Huang, Z. Li, Y. Wang, Z. Deng, J. Wang, P. C. Dedon, S. Chen, L. Wang, SspABCD–SspE is a phosphorothioation-sensing bacterial defence system with broad anti-phage activities. Nat. Microbiol., 1– 12 (2020).

5. A. Kuzmenko, A. Oguienko, D. Esyunina, D. Yudin, M. Petrova, A. Kudinova, O. Maslova, M. Ninova, S. Ryazansky, D. Leach, A. A. Aravin, A. Kulbachinskiy, DNA targeting and interference by a bacterial Argonaute nuclease. Nature, 1–10 (2020).

6. D. Cohen, S. Melamed, A. Millman, G. Shulman, Y. Oppenheimer-Shaanan, A. Kacen, S. Doron, G. Amitai, R. Sorek, Cyclic GMP–AMP signalling protects bacteria against viral infection. Nature. 574, 1–5 (2019).

7. A. Millman, A. Bernheim, A. Stokar-Avihail, T. Fedorenko, M. Voichek, A. Leavitt, Y. Oppenheimer-Shaanan, R. Sorek, Bacterial Retrons Function In Anti-Phage Defense. Cell. 183, 1551-1561.e12 (2020).

8. J. Bobonis, K. Mitosch, A. Mateus, G. Kritikos, J. R. Elfenbein, M. M. Savitski, H. Andrews-Polymenis, A. Typas, Phage proteins block and trigger retron toxin/antitoxin systems. bioRxiv, (2020).

9. S. V. Owen, N. Wenner, C. L. Dulberger, E. V. Rodwell, A. Bowers-Barnard, N. Quinones-Olvera, D. J. Rigden, E. J. Rubin, E. C. Garner, M. Baym, J. C. D. Hinton, Prophage-encoded phage defence proteins with cognate self-immunity. bioRxiv, (2020).

10. F. Depardieu, J. P. Didier, A. Bernheim, A. Sherlock, H. Molina, B. Duclos, D. Bikard, A Eukaryotic-like Serine/Threonine Kinase Protects Staphylococci against Phages. Cell Host Microbe. 20, 471–481 (2016).

11. R. L. Dy, R. Przybilski, K. Semeijn, G. P. C. Salmond, P. C. Fineran, A widespread bacteriophage abortive infection system functions through a Type IV toxin-antitoxin mechanism. Nucleic Acids Res. 42, 4590–4605 (2014).

12. G. Ofir, E. Herbst, M. Baroz, D. Cohen, A. Millman, S. Doron, N. Tal, D. B. A. Malheiro, S. Malitsky, G. Amitai, R. Sorek, Antiviral activity of bacterial TIR domains via signaling molecules that trigger cell death. bioRxiv, (2021).

13. S. Kronheim, M. Daniel-Ivad, Z. Duan, S. Hwang, A. I. Wong, I. Mantel, J. R. Nodwell, K. L. Maxwell, A chemical defence against phage infection. Nature. 564, 283–286 (2018).

14. A. Bernheim, A. Millman, G. Ofir, G. Meitav, C. Avraham, H. Shomar, M. M. Rosenberg, N. Tal, S. Melamed, G. Amitai, R. Sorek, Prokaryotic viperins produce diverse antiviral molecules. Nature, 1–8 (2020).

15. K. S. Makarova, Y. I. Wolf, E. V. Koonin, Comparative genomics of defense systems in archaea and bacteria. Nucleic Acids Res. 41, 4360–4377 (2013).

16. K. S. Makarova, Y. I. Wolf, S. Snir, E. V. Koonin, Defense Islands in Bacterial and Archaeal Genomes and Prediction of Novel Defense Systems. J. Bacteriol. 193, 6039–6056 (2011).

17. S. Doron, S. Melamed, G. Ofir, A. Leavitt, A. Lopatina, M. Keren, G. Amitai, R. Sorek, Systematic discovery of antiphage defense systems in the microbial pangenome. Science. 359, eaar4120 (2018).

18. L. Gao, H. Altae-Tran, F. Böhning, K. S. Makarova, M. Segel, J. L. Schmid-Burgk, J. Koob, Y. I. Wolf, E. V Koonin, F. Zhang, Diverse enzymatic activities mediate antiviral immunity in prokaryotes. Science. 369, 1077–1084 (2020).

19. N. Cumby, A. R. Davidson, K. L. Maxwell, The moron comes of age. Bacteriophage. 2, e23146 (2012).

20. K. Schroven, A. Aertsen, R. Lavigne, Bacteriophages as drivers of bacterial virulence and their potential for biotechnological exploitation. FEMS Microbiol. Rev. 041, 1–15 (2020).

21. X. Wang, Y. Kim, Q. Ma, S. H. Hong, K. Pokusaeva, J. M. Sturino, T. K. Wood, Cryptic prophages help bacteria cope with adverse environments. Nat. Commun. 1, 1–9 (2010).

22. D. Debroas, C. Siguret, Viruses as key reservoirs of antibiotic resistance genes in the environment. ISME J.13, 2856–2867 (2019).

23. V. L. Taylor, A. D. Fitzpatrick, Z. Islam, K. L. Maxwell, The Diverse Impacts of Phage Morons on Bacterial Fitness and Virulence. Advances in Virus Research (Academic Press Inc., 2019), vol. 103, pp. 1–31

24. A. Uc-Mass, E. J. Loeza, M. De La Garza, G. Guarneros, J. Hernández-Sánchez, L. Kameyama, An orthologue of the cor gene is involved in the exclusion of temperate lambdoid phages. Evidence that Cor inactivates FhuA receptor functions. Virology. 329, 425–433 (2004).

25. S. McGrath, G. F. Fitzgerald, D. Van Sinderen, Identification and characterization of phage-resistance genes in temperate lactococcal bacteriophages. Mol. Microbiol. 43, 509–520 (2002).

26. J. Oberto, R. A. Weisberg, M. E. Gottesman, Structure and function of the nun gene and the immunity region of the lambdoid phage HK022. J. Mol. Biol. 207, 675–693 (1989).

27. L. Snyder, Phage-exclusion enzymes: a bonanza of biochemical and cell biology reagents? Mol. Microbiol. 15 (1995), pp. 415–420.

28. K. D. Seed, D. W. Lazinski, S. B. Calderwood, A. Camilli, A bacteriophage encodes its own CRISPR/Cas adaptive response to evade host innate immunity. Nature. 494, 489–491 (2013).

29. J. Bondy-Denomy, J. Qian, E. R. Westra, A. Buckling, D. S. Guttman, A. R. Davidson, K. L. Maxwell, Prophages mediate defense against phage infection through diverse mechanisms. ISME J. 10, 2854–2866 (2016).

30. R. M. Dedrick, D. Jacobs-Sera, C. A. Guerrero Bustamante, R. A. Garlena, T. N. Mavrich, W. H. Pope, J. C. Cervantes Reyes, D. A. Russell, T. Adair, R. Alvey, J. A. Bonilla, J. S. Bricker, B. R. Brown, D. Byrnes, S. G. Cresawn, W. B. Davis, L. A. Dickson, N. P. Edgington, A. M. Findley, U. Golebiewska, J. H. Grose, C. F. Hayes, L. E. Hughes, K. W. Hutchison, S. Isern, A. A. Johnson, M. A. Kenna, K. K. Klyczek, C. M. Mageeney, S. F. Michael, S. D. Molloy, M. T. Montgomery, J. Neitzel, S. T. Page, M. C. Pizzorno, M. K. Poxleitner, C. A. Rinehart, C. J. Robinson, M. R. Rubin, J. N. Teyim, E. Vazquez, V. C. Ware, J. Washington, G. F. Hatfull, Prophage-mediated defence against viral attack and viral counter-defence. Nat. Microbiol. 2, 1–13 (2017).

31. A. Fillol-Salom, L. Miguel-Romero, A. Marina, J. Chen, J. R. Penadés, Beyond the CRISPR-Cas safeguard: PICI-encoded innate immune systems protect bacteria from bacteriophage predation. Curr. Opin. Microbiol. 56, 52–58 (2020).

32. E. V. Koonin, K. S. Makarova, Y. I. Wolf, M. Krupovic, Evolutionary entanglement of mobile genetic elements and host defence systems: guns for hire. Nat. Rev. Genet. (2019),, doi:10.1038/s41576-019-0172-9.

33. A. S. Nilsson, J. L. Karlsson, E. Haggård-Ljungquist, Site-Specific Recombination Links the Evolution of P2-like Coliphages and Pathogenic Enterobacteria. Mol. Biol. Evol. 21, 1–13 (2004).

34. R. Pinilla-Redondo, S. Shehreen, N. D. Marino, R. D. Fagerlund, C. M. Brown, S. J. Sørensen, P. C. Fineran, J. Bondy-Denomy, Discovery of multiple anti-CRISPRs highlights anti-defense gene clustering in mobile genetic elements. Nat. Commun. 11, 5652 (2020).

35. A. Pawluk, J. Bondy-Denomy, V. H. W. Cheung, K. L. Maxwell, A. R. Davidson, A new group of phage anti-CRISPR genes inhibits the type I-E CRISPR-Cas system of pseudomonas aeruginosa. MBio. 5 (2014), doi:10.1128/mBio.00896-14.

36. F. Rousset, J. Cabezas-Caballero, F. Piastra-Facon, O. Clermont, E. Denamur, E. P. C Rocha, D. Bikard, The impact of genetic diversity on gene essentiality within the Escherichia coli species. Nat. Microbiol., 1–12 (2021).

37. J. Bobonis, A. Mateus, B. Pfalz, S. Garcia-Santamarina, M. Galardini, C. Kobayashi, F. Stein, M. M. Savitski, J. R. Elfenbein, H. Andrews-Poymenis, A. Typas, Bacterial retrons encode tripartite toxin/antitoxin systems. bioRxiv. (2020).

38. L. M. Bobay, M. Touchon, E. P. C. Rocha, Pervasive domestication of defective prophages by bacteria. Proc. Natl. Acad. Sci. U. S. A. 111, 12127–12132 (2014).

39. K. Kita, J. Tsuda, T. Kato, K. Okamoto, H. Yanase, M. Tanaka, Evidence of horizontal transfer of the EcoO109I restriction-modification gene to Escherichia coli chromosomal DNA. J. Bacteriol. 181, 6822–6827 (1999).

40. S. Inouye, M. G. Sunshine, E. W. Six, M. Inouye, Retronphage φR73: An E. coli phage that contains a retroelement and integrates into a tRNA gene. Science (80-.). 252, 969–971 (1991).

41. E. Koyuncu, H. G. Budayeva, Y. V. Miteva, D. P. Ricci, T. J. Silhavy, T. Shenk, I. M. Cristea, Sirtuins are evolutionarily conserved viral restriction factors. MBio. 5 (2014), doi:10.1128/mBio.02249-14.

42. V. Anantharaman, K. S. Makarova, A. M. Burroughs, E. V. Koonin, L. Aravind, Comprehensive analysis of the HEPN superfamily: Identification of novel roles in intra-genomic conflicts, defense, pathogenesis and RNA processing. Biol. Direct. 8, 1–28 (2013).

43. K. Takeda, S. Akira, Toll-like receptors in innate immunity. Int. Immunol. 17 (2005), pp. 1–14.

44. E. V. Koonin, L. Aravind, Origin and evolution of eukaryotic apoptosis: The bacterial connection. Cell Death Differ. 9 (2002), pp. 394–404.

45. A. Lopatina, N. Tal, R. Sorek, Abortive Infection: Bacterial Suicide as an Antiviral Immune Strategy. Annu. Rev. Virol., (2020).

46. F. W. Studier, Gene 0.3 of bacteriophage T7 acts to overcome the DNA restriction system of the host. J. Mol. Biol. 94, 283–295 (1975).

47. A. Isaev, A. Drobiazko, N. Sierro, J. Gordeeva, I. Yosef, U. Qimron, N. V Ivanov, K. Severinov, Phage T7 DNA mimic protein Ocr is a potent inhibitor of BREX defence. Nucleic Acids Res. 48, 5397–5406 (2020).

48. M. Penner, I. Morad, L. Snyder, G. Kaufmann, Phage T4-coded Stp: Double-edged effector of coupled DNA and tRNA-restriction systems. J. Mol. Biol. 249, 857–868 (1995).

49. A. J. Enright, I. Illopoulos, N. C. Kyrpides, C. A. Ouzounis, Protein interaction maps for complete genomes based on gene fusion events. Nature. 402, 86–90 (1999).

50. B. H. Lindqvist, G. Deho, R. Calendar, Mechanisms of genome propagation and helper exploitation by satellite phage P4. Microbiol. Rev. 57 (1993), pp. 683–702.

51. R. Ziermann, R. Calendar, Characterization of the cos sites of bacteriophages P2 and P4. Gene. 96, 9–15 (1990).

52. G. E. Christie, R. Calendar, Bacteriophage P2. Bacteriophage. 6, e1145782 (2016).

53. W. M. Johnson, H. Lior, A new heat-labile cytolethal distending toxin (CLDT) produced by Escherichia coli isolates from clinical material. Microb. Pathog. 4, 103–113 (1988).

54. D. Shore, G. Deho, J. Tsipis, R. Goldstein, Determination of capsid size by satellite bacteriophage P4. Proc. Natl. Acad. Sci. U. S. A. 75, 400–404 (1978).

55. I. Mortier-Barrière, M. Velten, P. Dupaigne, N. Mirouze, O. Piétrement, S. McGovern, G. Fichant, B. Martin, P. Noirot, E. Le Cam, P. Polard, J. P. Claverys, A Key Presynaptic Role in Transformation for a Widespread Bacterial Protein: DprA Conveys Incoming ssDNA to RecA. Cell. 130, 824–836 (2007).

56. G. R. Dwivedi, E. Sharma, D. N. Rao, Helicobacter pylori DprA alleviates restriction barrier for incoming DNA. Nucleic Acids Res. 41, 3274–3288 (2013).

57. M. R. Mestre, A. González-Delgado, L. I. Gutíerrez-Rus, F. Martínez-Abarca, N. Toro, Systematic prediction of genes functionally associated with bacterial retrons and classification of the encoded tripartite systems. Nucleic Acids Res. 48, 12632–12647 (2020).

58. D. Kapfhammer, J. Blass, S. Evers, J. Reidl, Vibrio cholerae phage K139: Complete genome sequence and comparative genomics of related phages. J. Bacteriol. 184, 6592–6601 (2002).

59. F. Beilstein, B. Dreiseikelmann, Temperate bacteriophage φO18P from an Aeromonas media isolate: Characterization and complete genome sequence. Virology. 373, 25–29 (2008).

60. K. N. LeGault, S. G. Hays, A. Angermeyer, A. C. McKitterick, F. Johura, M. Sultana, M. Alam, K. D. Seed, Temporal Shifts in Antibiotic Resistance Elements Govern Virus-Pathogen Conflicts. bioRxiv (2020).

61. C. M. Johnson, M. M. Harden, A. D. Grossman, An integrative and conjugative element encodes an abortive infection system to protect host cells from predation by a bacteriophage. bioRxiv (2020).

62. P. H. Oliveira, M. Touchon, J. Cury, E. P. C. Rocha, The chromosomal organization of horizontal gene transfer in bacteria. Nat. Commun. 8 (2017), doi:10.1038/s41467-017-00808-w.

63. M. Touchon, L. M. Bobay, E. P. C. Rocha, The chromosomal accommodation and domestication of mobile genetic elements. Curr. Opin. Microbiol. 22 (2014), pp. 22–29.

64. M. Steinegger, J. Söding, MMseqs2 enables sensitive protein sequence searching for the analysis of massive data sets. Nat. Biotechnol. 35, 1026–1028 (2017).

65. C. L. M. Gilchrist, Y.-H. Chooi, clinker & clustermap.js: Automatic generation of gene cluster comparison figures. bioRxiv (2020).

66. S. El-Gebali, J. Mistry, A. Bateman, S. R. Eddy, A. Luciani, S. C. Potter, M. Qureshi, L. J. Richardson, G. A. Salazar, A. Smart, E. L. L. Sonnhammer, L. Hirsh, L. Paladin, D. Piovesan, S. C. E. Tosatto, R. D. Finn, The Pfam protein families database in 2019. Nucleic Acids Res. 47, D427–D432 (2019).

67. S. S. Abby, B. Néron, H. Ménager, M. Touchon, E. P. C. Rocha, MacSyFinder: A Program to Mine Genomes for Molecular Systems with an Application to CRISPR-Cas Systems. PLoS One. 9, e110726 (2014).

68. D. G. Gibson, L. Young, R.-Y. Chuang, J. C. Venter, C. A. Hutchison, H. O. Smith, Enzymatic assembly of DNA molecules up to several hundred kilobases. Nat. Methods. 6, 343–345 (2009).

69. D. Maura, E. Morello, L. du Merle, P. Bomme, C. L. Bouguénec, L. Debarbieux, Intestinal colonization by enteroaggregative Escherichia coli supports long-term bacteriophage replication in mice. Environ. Microbiol. 14, 1844–1854 (2012).

70. M. Galtier, L. De Sordi, A. Sivignon, A. de Vallée, D. Maura, C. Neut, O. Rahmouni, K. Wannerberger, A. Darfeuille-Michaud, P. Desreumaux, N. Barnich, L. Debarbieux, Bacteriophages targeting adherent invasive Escherichia coli strains as a promising new treatment for Crohn’s disease. J. Crohn’s Colitis. 11, 840–847 (2017).

71. M. Galtier, L. De Sordi, D. Maura, H. Arachchi, S. Volant, M. Dillies, L. Debarbieux, Bacteriophages to reduce gut carriage of antibiotic resistant uropathogens with low impact on microbiota composition. Environ. Microbiol. 18, 2237–2245 (2016).

72. I. M. A. Chen, K. Chu, K. Palaniappan, M. Pillay, A. Ratner, J. Huang, M. Huntemann, N. Varghese, J. R. White, R. Seshadri, T. Smirnova, E. Kirton, S. P. Jungbluth, T. Woyke, E. A. Eloe-Fadrosh, N. N. Ivanova, N. C. Kyrpides, IMG/M v.5.0: An integrated data management and comparative analysis system for microbial genomes and microbiomes. Nucleic Acids Res. 47, D666–D677 (2019).

