## Supplementary Figures for "Prophage-encoded hotspots of bacterial immune systems"

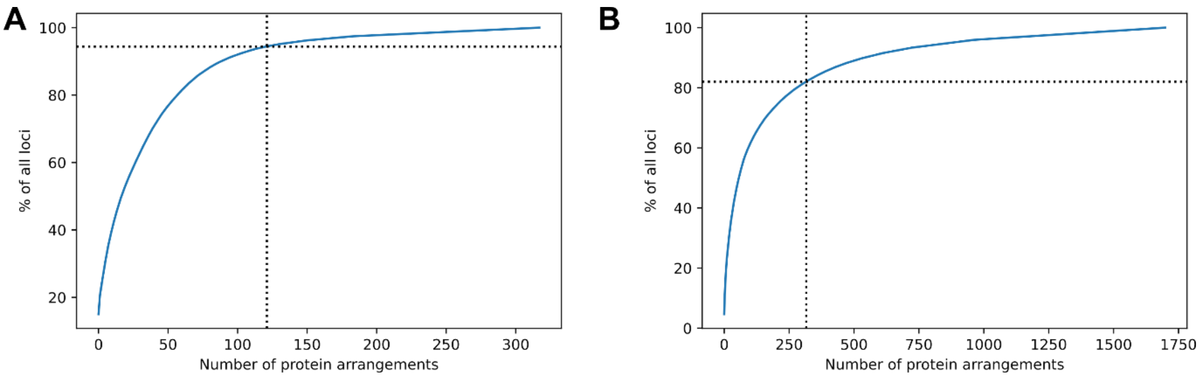


**Fig. S1. Distribution of the relative abundance of gene arrangements.** Plots show the evolution of the frequency of each gene arrangement from P4-encoded (**A**) or P2-encoded (**B**) hotspots. Dashed lines highlight the analyzed threshold: (A) 121 arrangements occurring at least 5 times in the P4-encoded hostpot account for 94.4% of all P4-encoded loci, while (B) 316 arrangements occurring at least 10 times in the P2-encoded hotspot account for 82% of all P2-encoded loci.

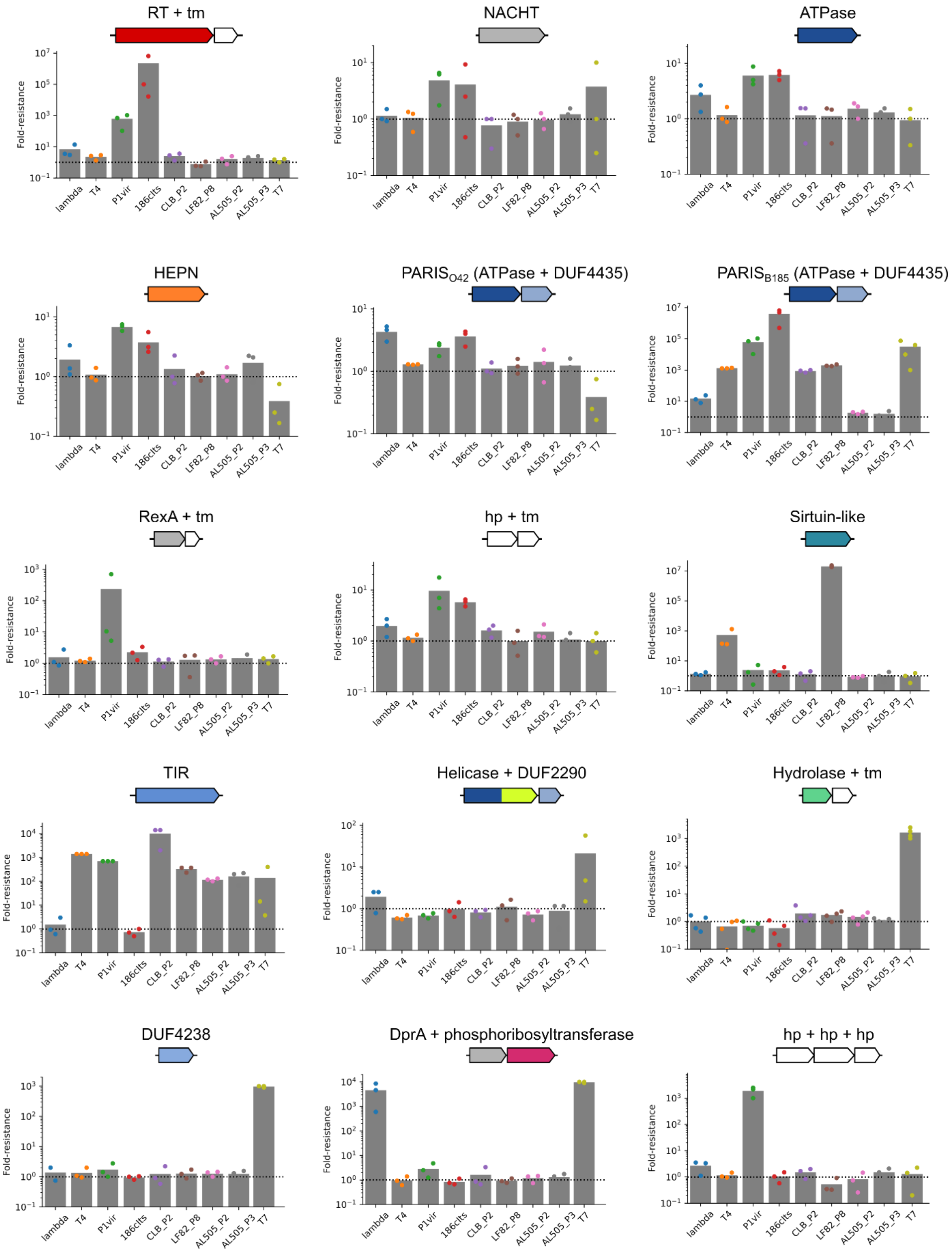


**Fig. S2. Detailed fold-resistance of all verified defense systems against 9 phages (related to Fig. 2).** The number of plaques was measured for each of 9 phages on cells harboring either a control plasmid or a putative defense system. Fold-resistance was measured as the ratio between these two values. Bar plots show the mean of 3 to 4 independent measurements. RT: reverse-transcriptase, tm; transmembrane domain, DUF: domain of unknown function, hp: hypothetical protein, TIR: Toll/Interleukin-1 Receptor

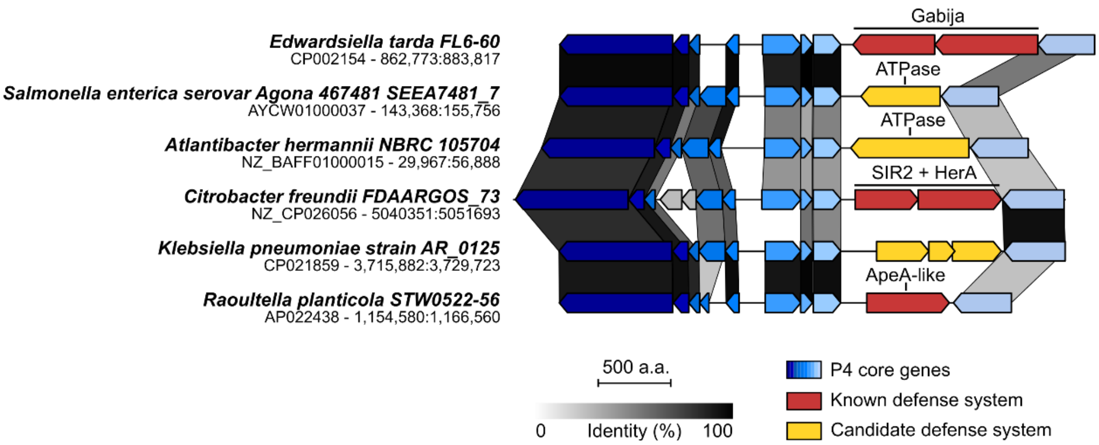


**Fig. S3. Genetic diversity encoded on P4-like phages outside *E. coli*.** Genomic comparison of the P4-like prophage encoded in six different genomes of Enterobacteriaceae. P4 core genes are shown in shades of blue. Grey shades show the percentage of identity between homologous proteins from different genomes. Genome accession numbers and positions are shown on the left.

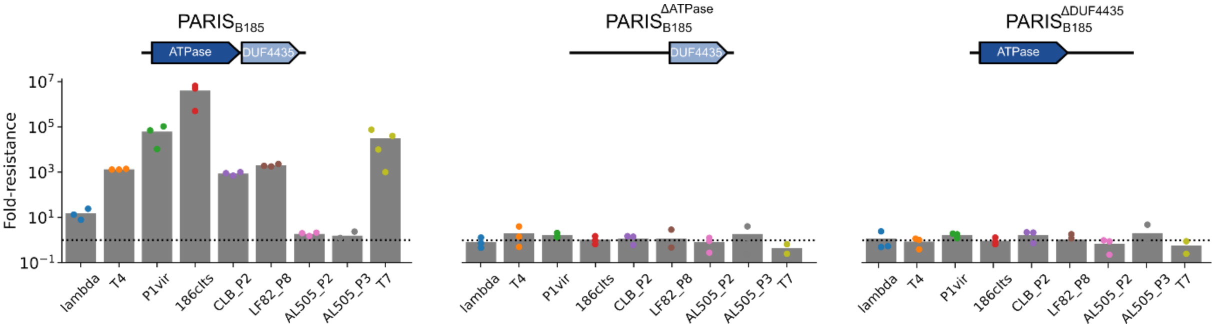


**Fig. S4. Deletion of either protein from PARIS abrogates defense.** Fold-resistance against our phage panel was measured for the wild-type PARIS from *E. coli* B185 (left) or for deletion mutants of the ATPase (middle) or the DUF4435 protein (right) (**see Methods**). Bar plots show the mean of 3 independent measurements.

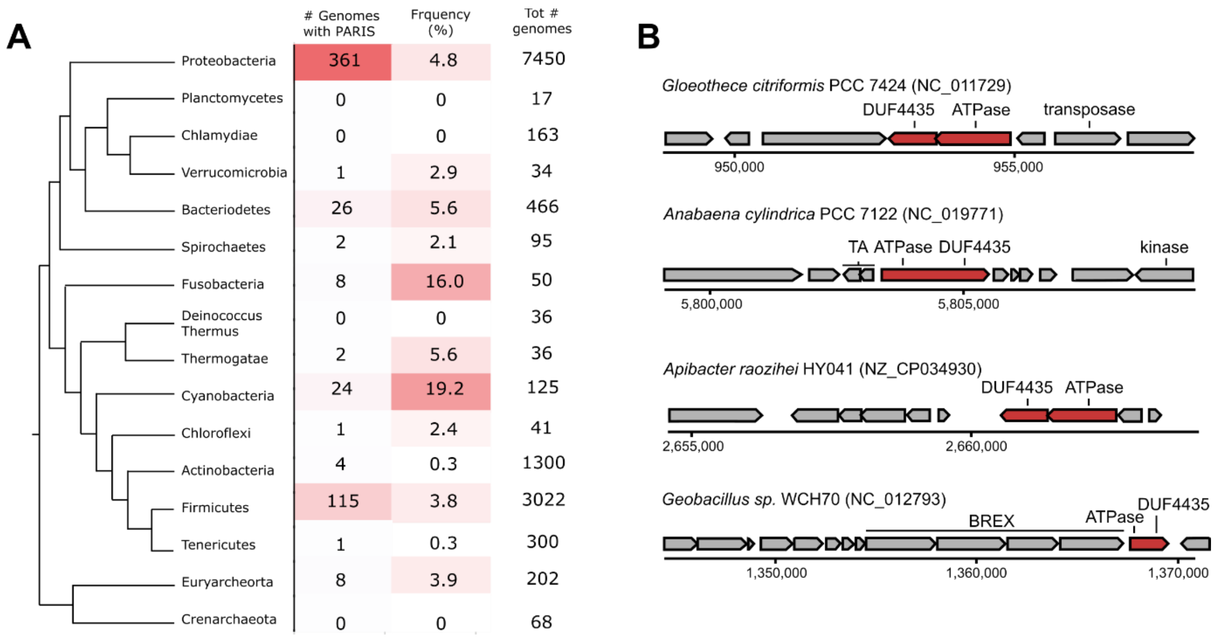


**Fig. S5. Detection of PARIS in Bacteria and Archaea.** Systems were searched in 13,512 prokaryotic genomes using MacSyFinder (see Methods). (**A**) For each clade, cells show the number and proportion of genomes where we detected PARIS. A color gradient is used to depict the prevalence. The cladogram on the left shows the approximate phylogenetic relationship between clades. (**B**) Genomic view of a few occurrences of PARIS in bacterial genomes. PARIS occurs either as a two-protein system or as a single protein fusion. PARIS genes are colored in red. Genome accessions are shown in parentheses and the bottom tracks show genome coordinates.

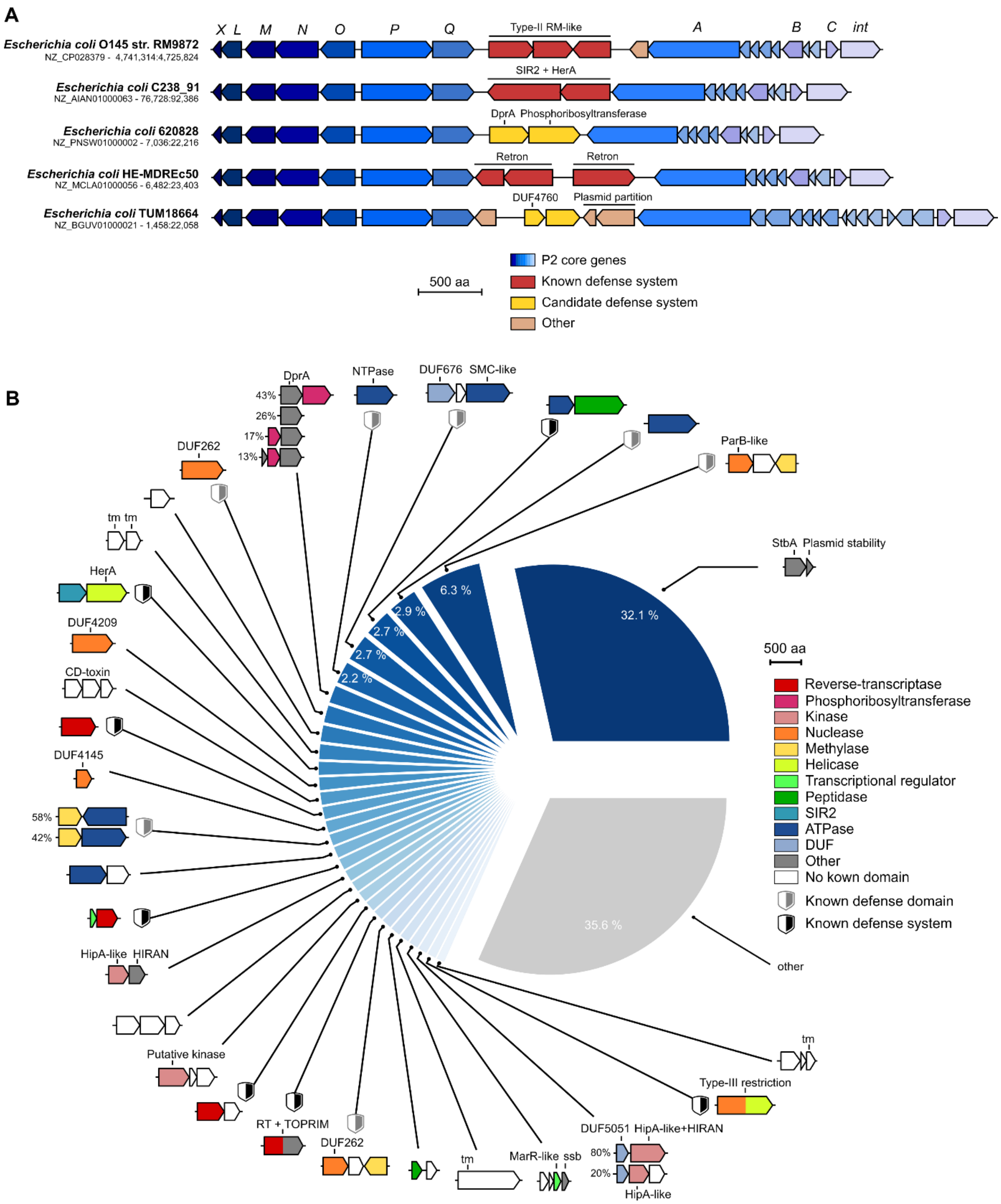


**Fig. S6. A diversity of genetic systems encoded on P2-like phages in *E. coli*.** (**A**) Visualization of genomic regions from five *E. coli* strains containing a P2-like prophage, highlighting genetic diversity between *Q* and *A* genes, including known anti-phage defense systems. Genome accession numbers and positions are shown on the left. (**B**) Systematic analysis of genetic systems encoded between gpA and gpQ from P2-like phages. The pie chart shows the 30 most abundant systems classified by prevalence and shown as gene cassettes colored by protein domains (not to scale). The percentage of loci encoding each system is shown for systems present in at least 2% of loci. Note that most loci encode more than one system, so that the sum is greater than 100%. When a system comprises accessory genes, different variants are shown with the percentage of each occurrence shown on the left. TIR: Toll/Interleukin-1 Receptor, HAD: haloacid dehydrogenase-like, SIR2: sirtuin, DUF: domain of unknown function, tm: transmembrane domain, SMC: Structural Maintenance of Chromosome.

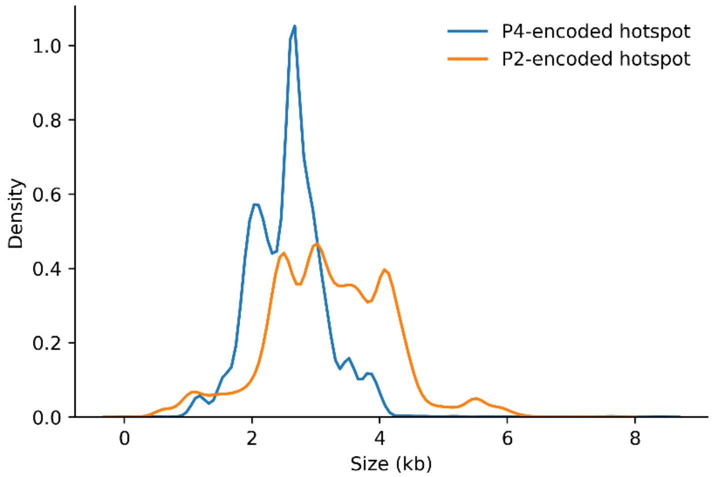


**Fig. S7. Distribution of the size of P2- and P4-encoded hotspots**. We measured the size of the P4-encoded hotspot at each locus as the genomic distance between the end of *psu* and the end of *int*. For the P2-encoded hotspot, we used the distance between the end of *A* and the end of *Q*. P2-like phages encode larger systems than P4-like phages (Mann-Whitney p-value $< 10^{-100}$).

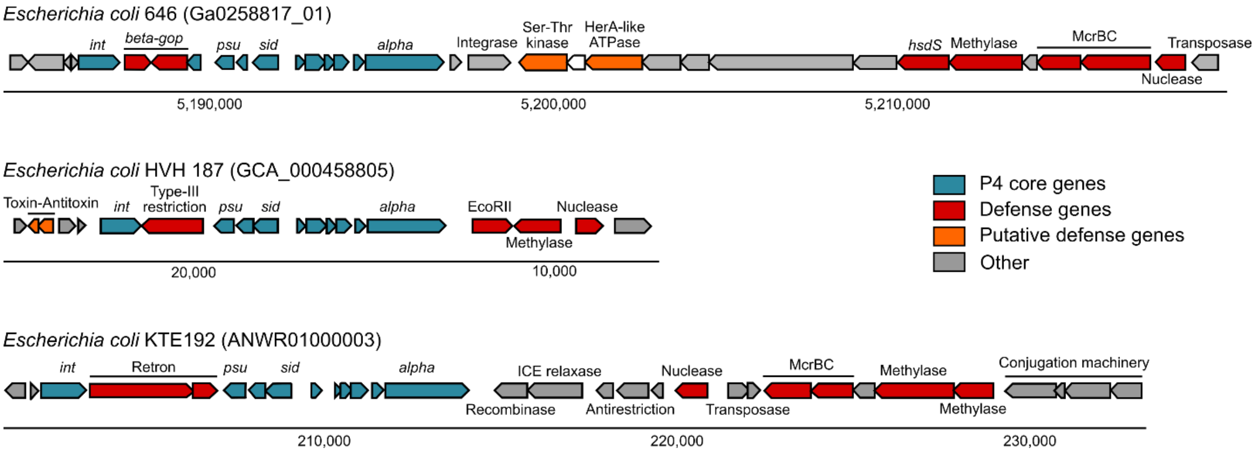


**Fig. S8. Genomic view of P4-like elements integrated near defense systems and integrative conjugative elements.** P4 genes are colored in blue while defense-associated genes are colored in orange or red. Genome accession numbers are shown in parentheses and the bottom tracks shows genomic coordinates.

**Table S1. Accessions of all P4-encoded loci.** This table lists all proteins from P4-encoded hotspots detected in all *E. coli* genomes. The “Genome accession” column provides the RefSeq genome ID. The “Pfams” column refers to the detected protein families within the hotspot proteins. The “Protein accessions” column provides the RefSeq protein ID of the hotspot-encoded proteins.

**Table S2. Summary of genetic systems encoded on P4-like satellites.** This table summarizes all curated genetic systems encoded in P4 hotspots with at least 5 occurrences.

**Table S3. List of phages used in this study.**

**Table S4. Description of validated defense systems.** All experimentally validated systems (Fig. 2) are detailed with the source (“Source”, “Genome accession” and “Locus”), cloning information (“Cloned sequence”, “Forward primer” and “Reverse primer”) and content (“Number of genes” and protein sequences).

**Table S5. Accessions of all P2-encoded loci.** This table lists all proteins from P2-encoded hotspots detected in all *E. coli* genomes. The “Genome accession” column provides the RefSeq genome ID. The “Pfams” column refers to the detected protein families within the hotspot proteins. The “Protein accessions” column provides the RefSeq protein ID of the hotspot-encoded proteins.

**Table S6. DNA sequence of the plasmid used to test defense systems**
